## Supplementary material for "A comparative study of supervised machine learning algorithms for the prediction of long-range chromatin interactions"

### Figure S1

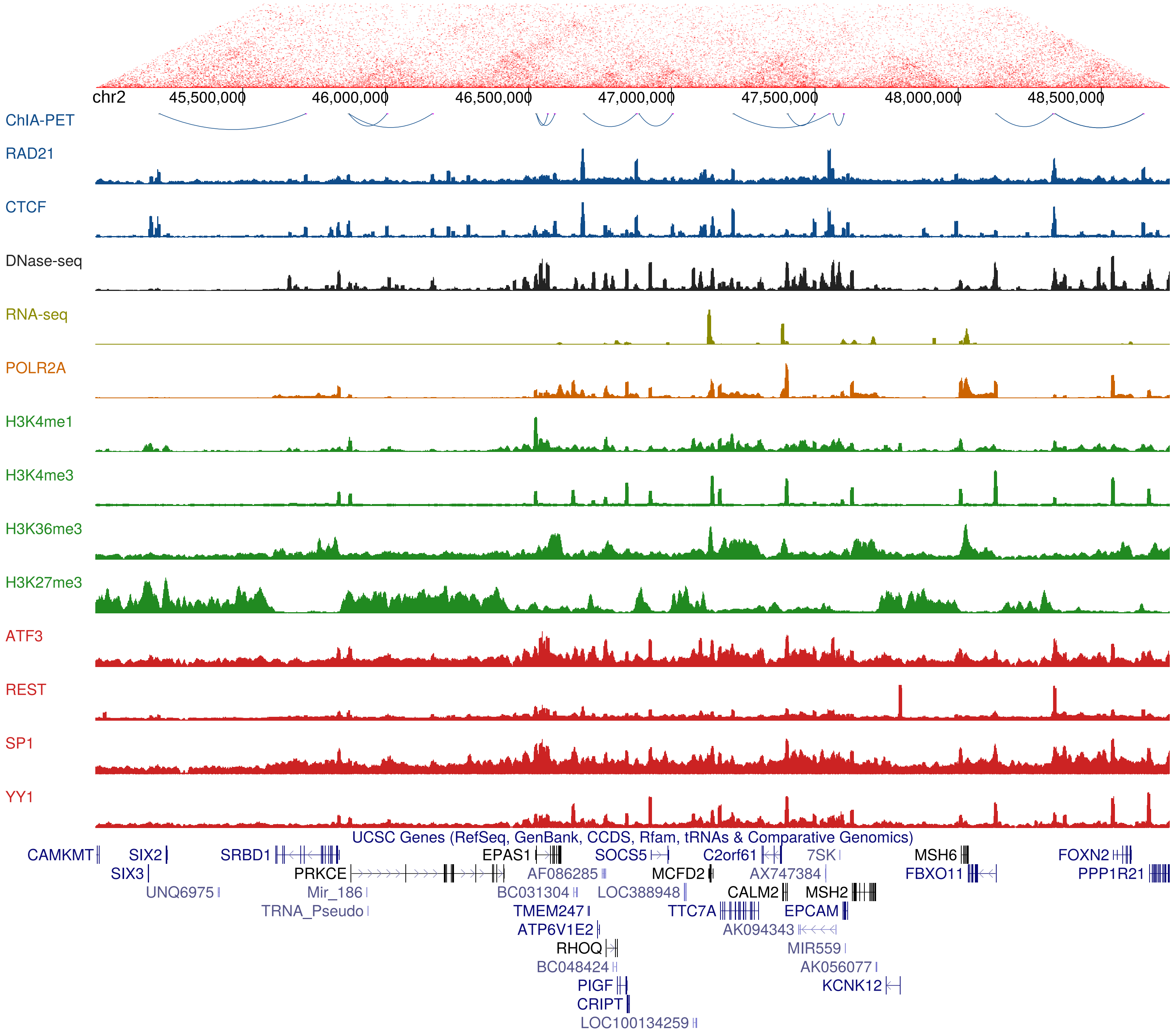
chr2
45,500,000
46,000,000
46,500,000
47,000,000
47,500,000
48,000,000
48,500,000
ChIA-PET
RAD21
CTCF
DNase-seq
RNA-seq
POLR2A
H3K4me1
H3K4me3
H3K36me3
H3K27me3
ATF3
REST
SP1
YY1
UCSC Genes (RefSeq, GenBank, CCDS, Rfam, tRNAs & Comparative Genomics)
CAMKMT
SIX2
SRBD1
EPAS1
SOCS5
C2orf61
7SK
MSH6
FOXN2
SIX3
PRKCE
AF086285
MCFD2
AX747384
FBXO11
PPP1R21
UNQ6975
Mir_186
BC031304
LOC388948
CALM2
MSH2
TRNA_Pseudo
TMEM247
TTC7A
EPCAM
ATP6V1E2
AK094343
RHOQ
MIR559
BC048424
AK056077
PIGF
KCNK12
CRIPT
LOC100134259

### Figure S2

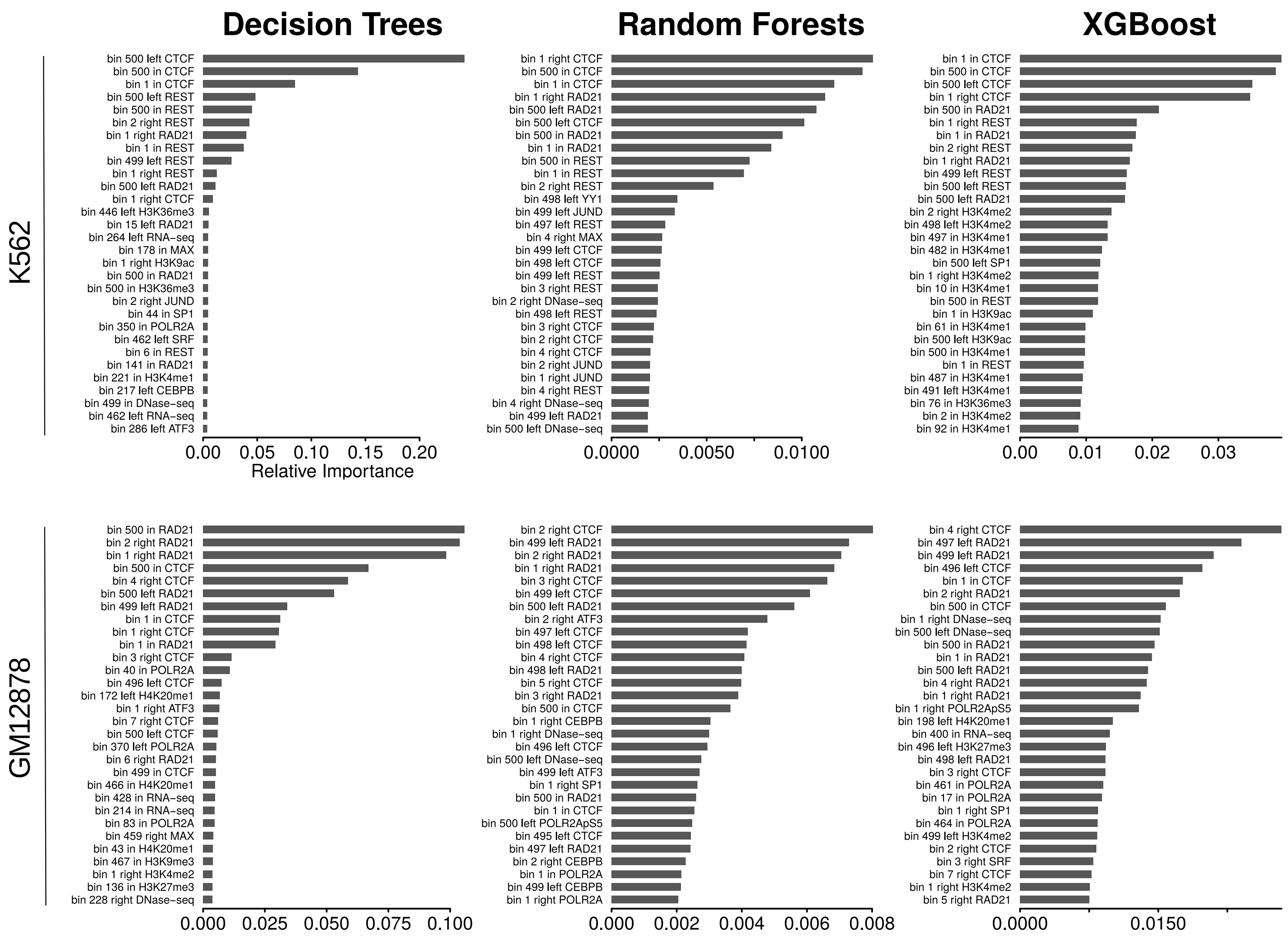
Decision Trees
Random Forests
XGBoost
K562
GM12878
Relative Importance

#### **Figure S3**

#### **A)** Decision Trees (K562)

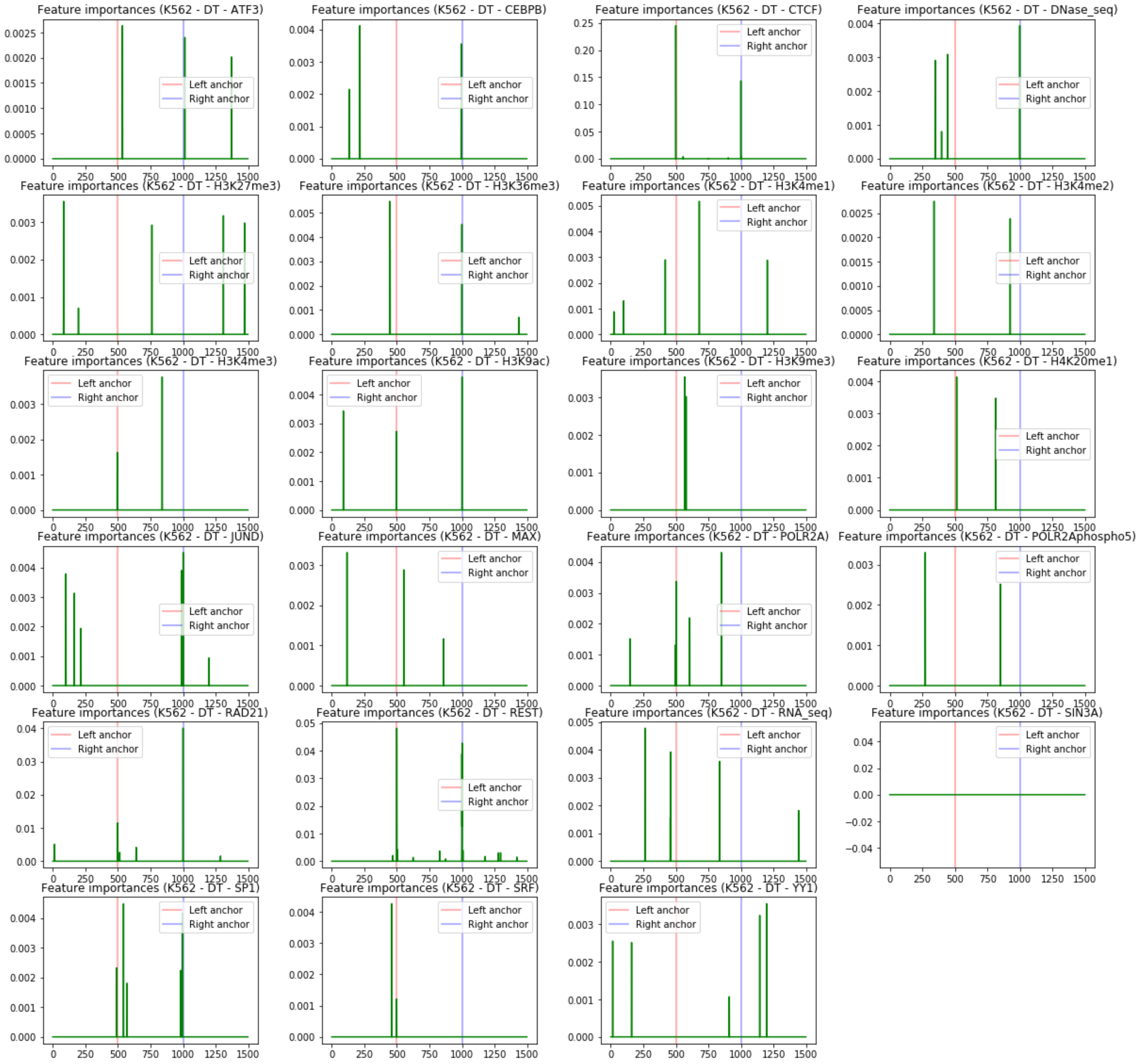

**B)** Random Forests (K562)

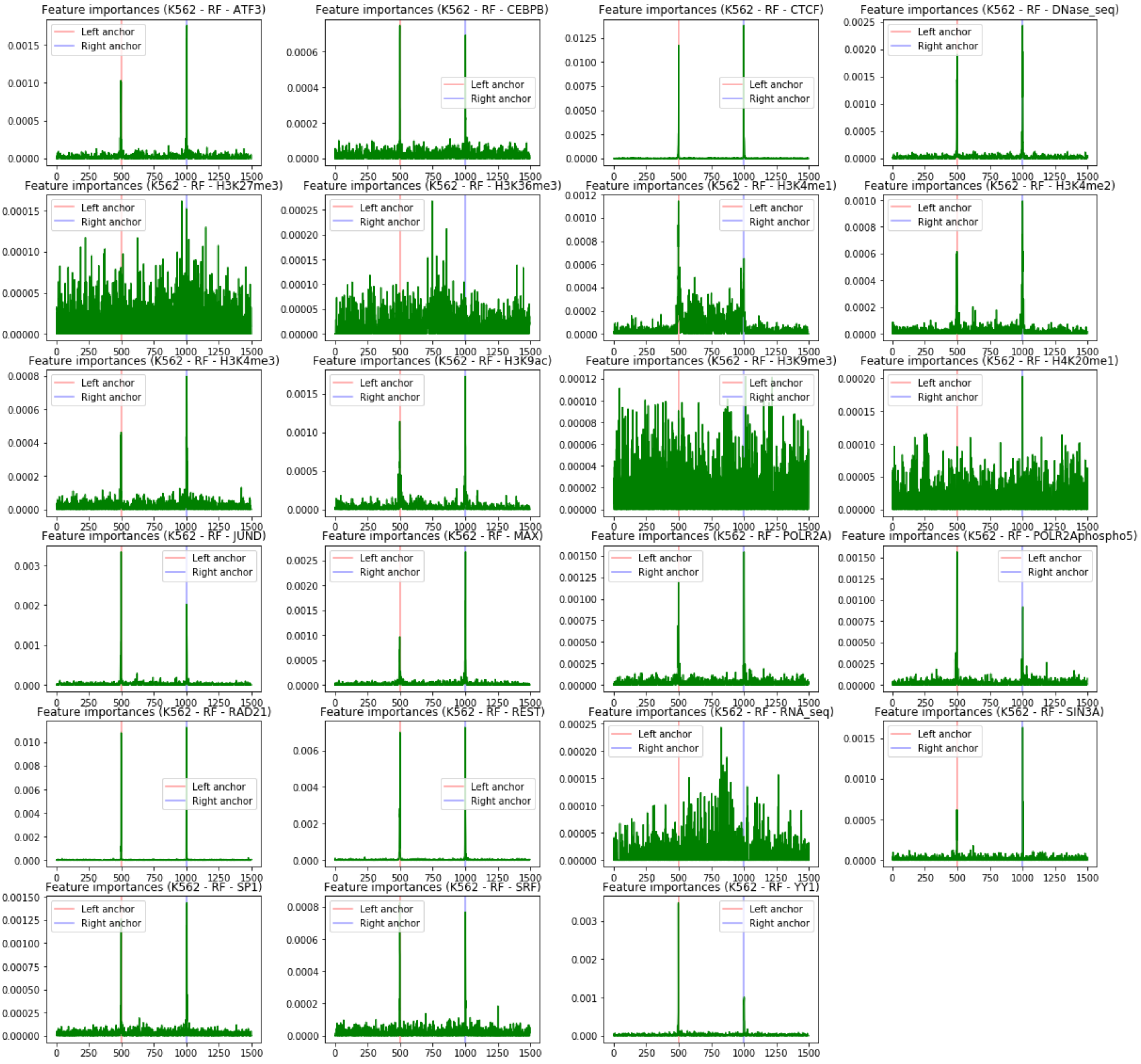

#### **C)** XGBoost (K562)

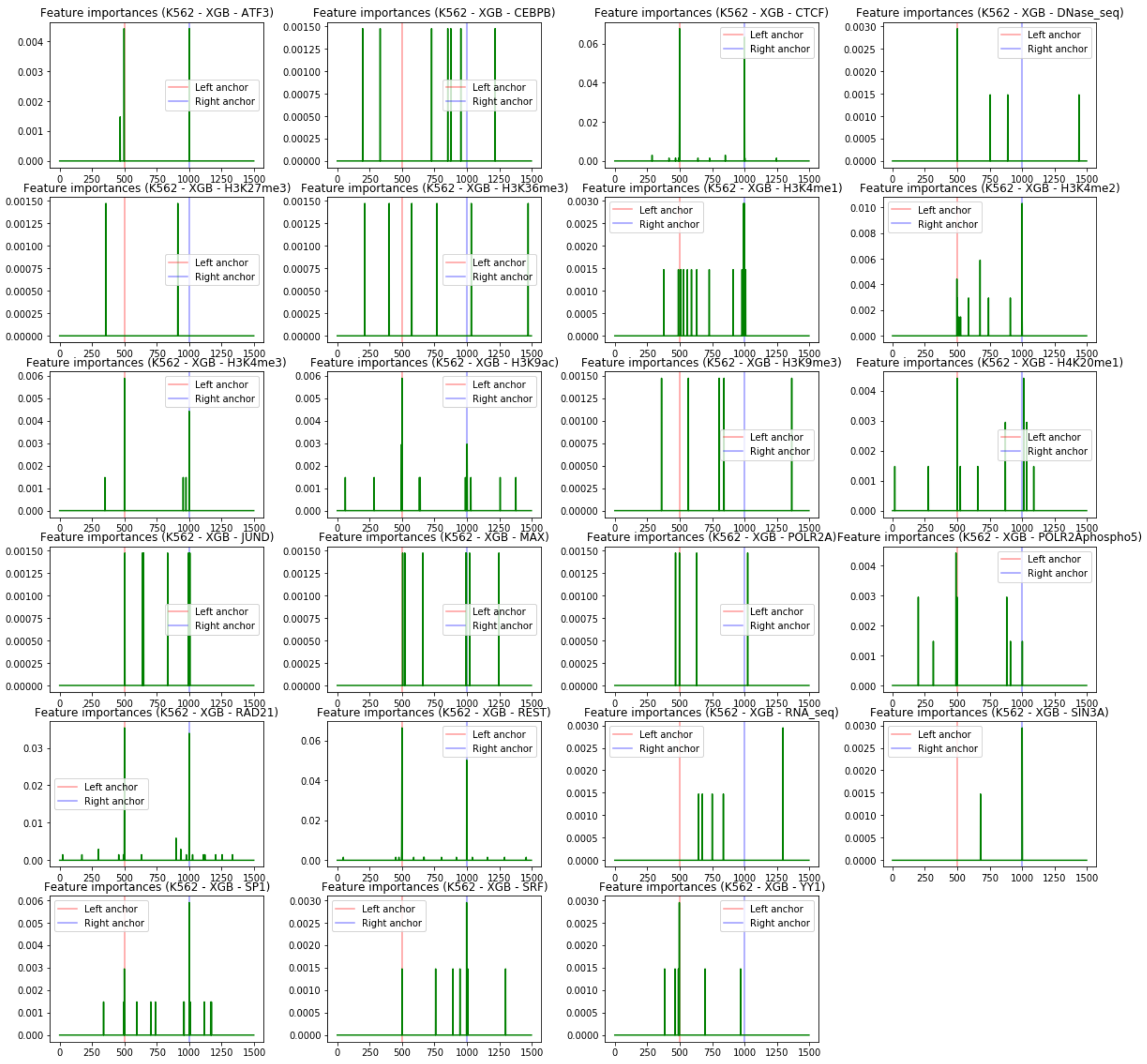

#### **D)** Decision Trees (GM12878)

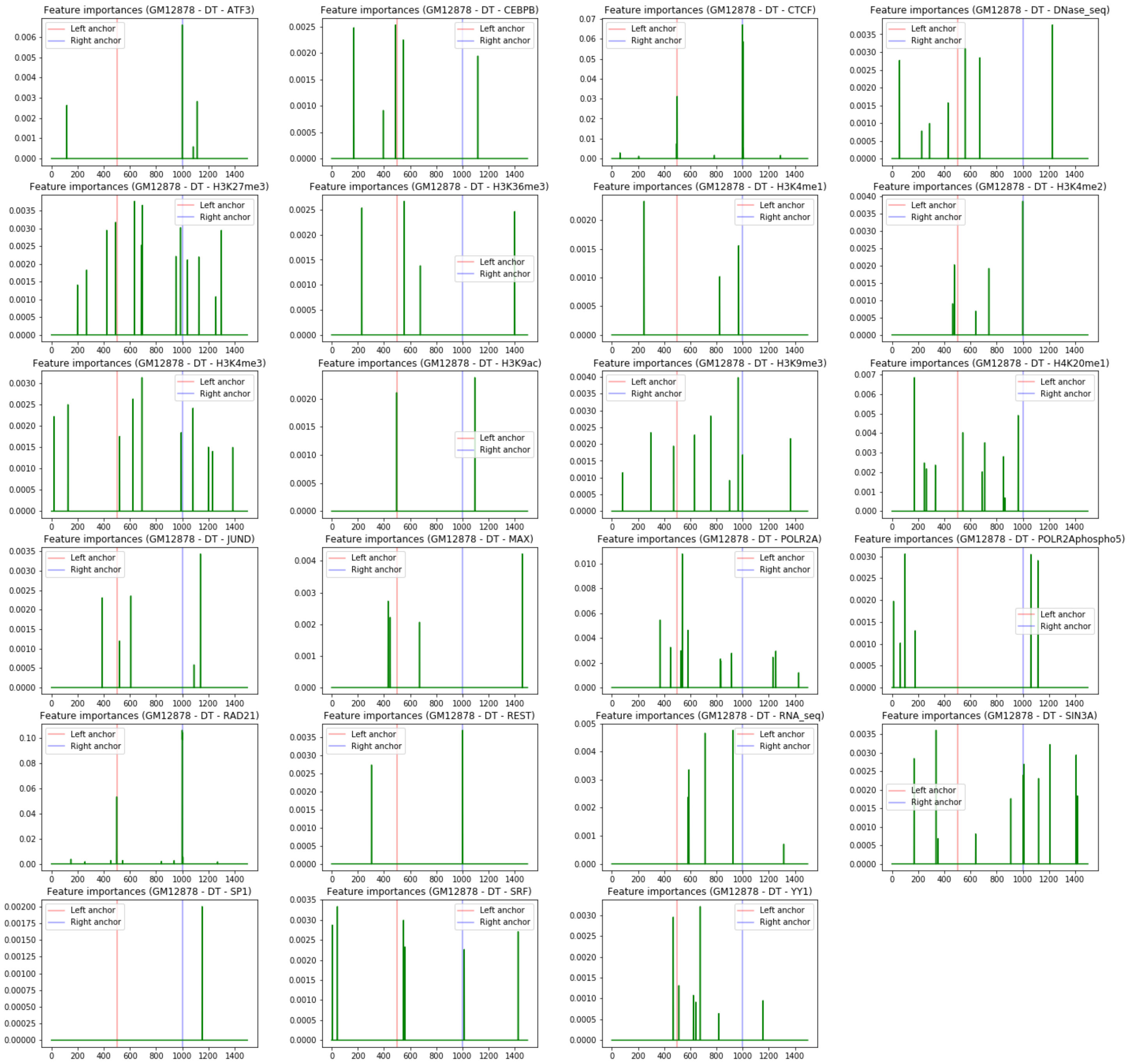

**E)** Random Forests (GM12878)

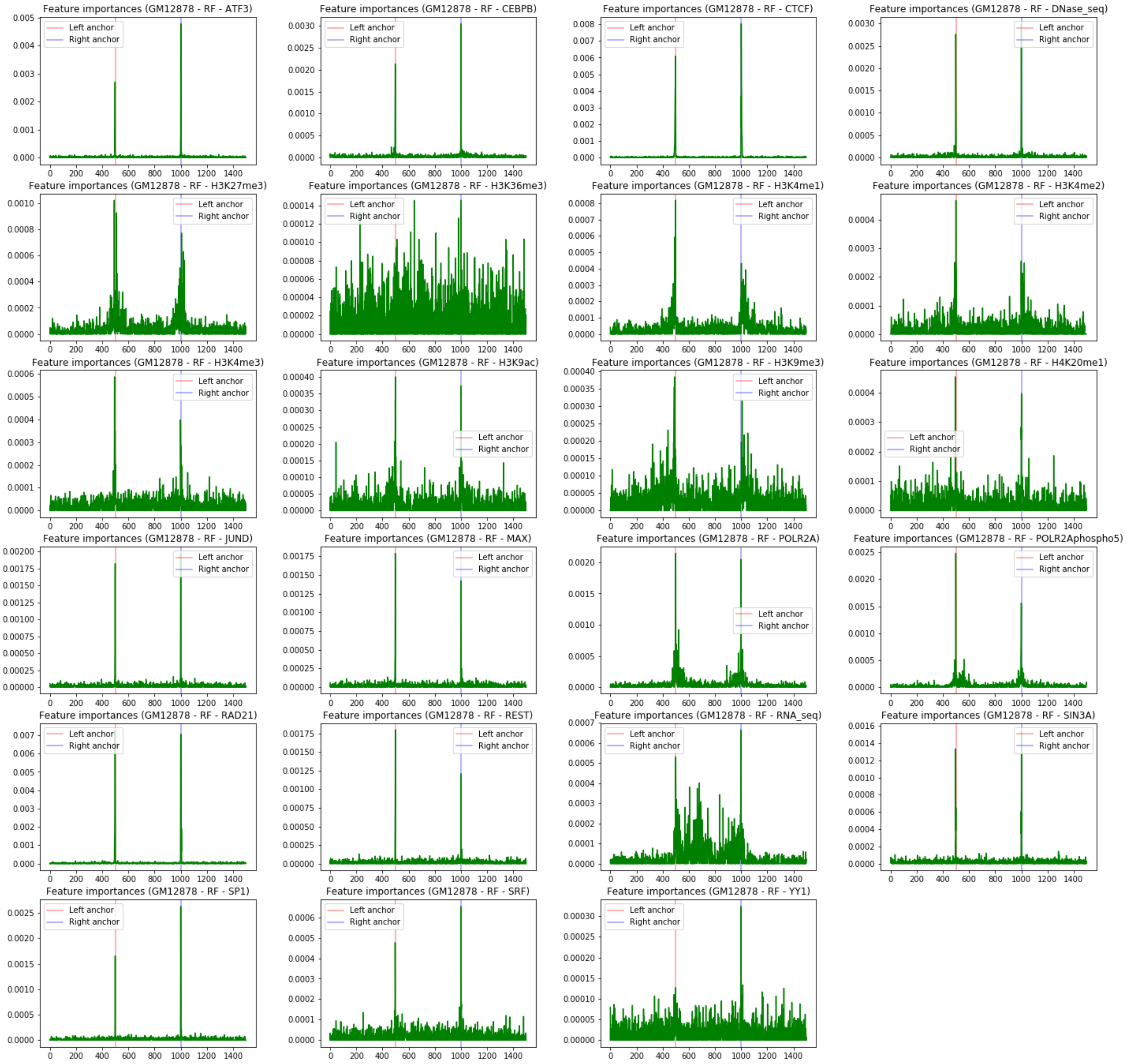

**F)** XGBoost (GM12878)

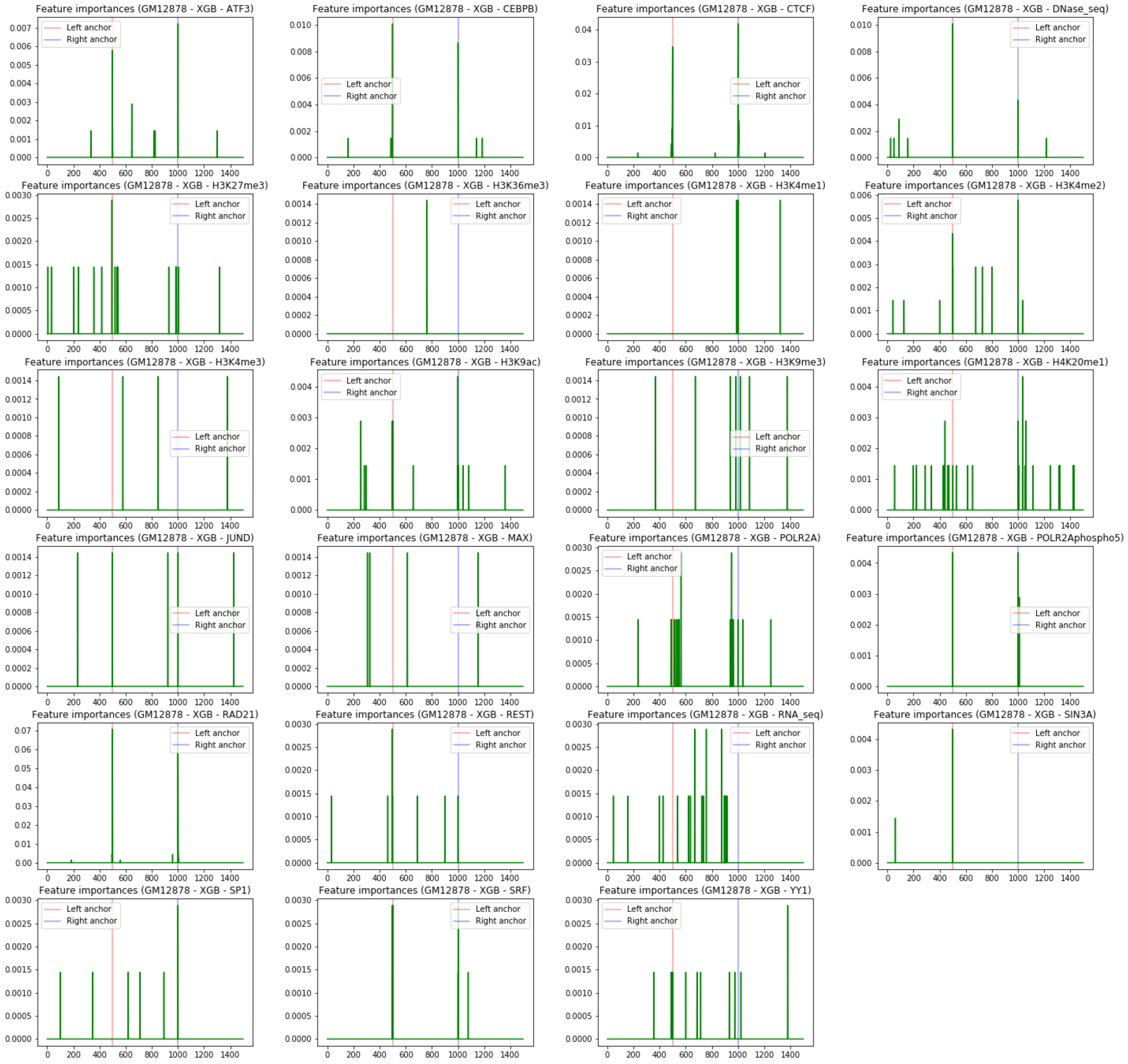

#### Figure S4

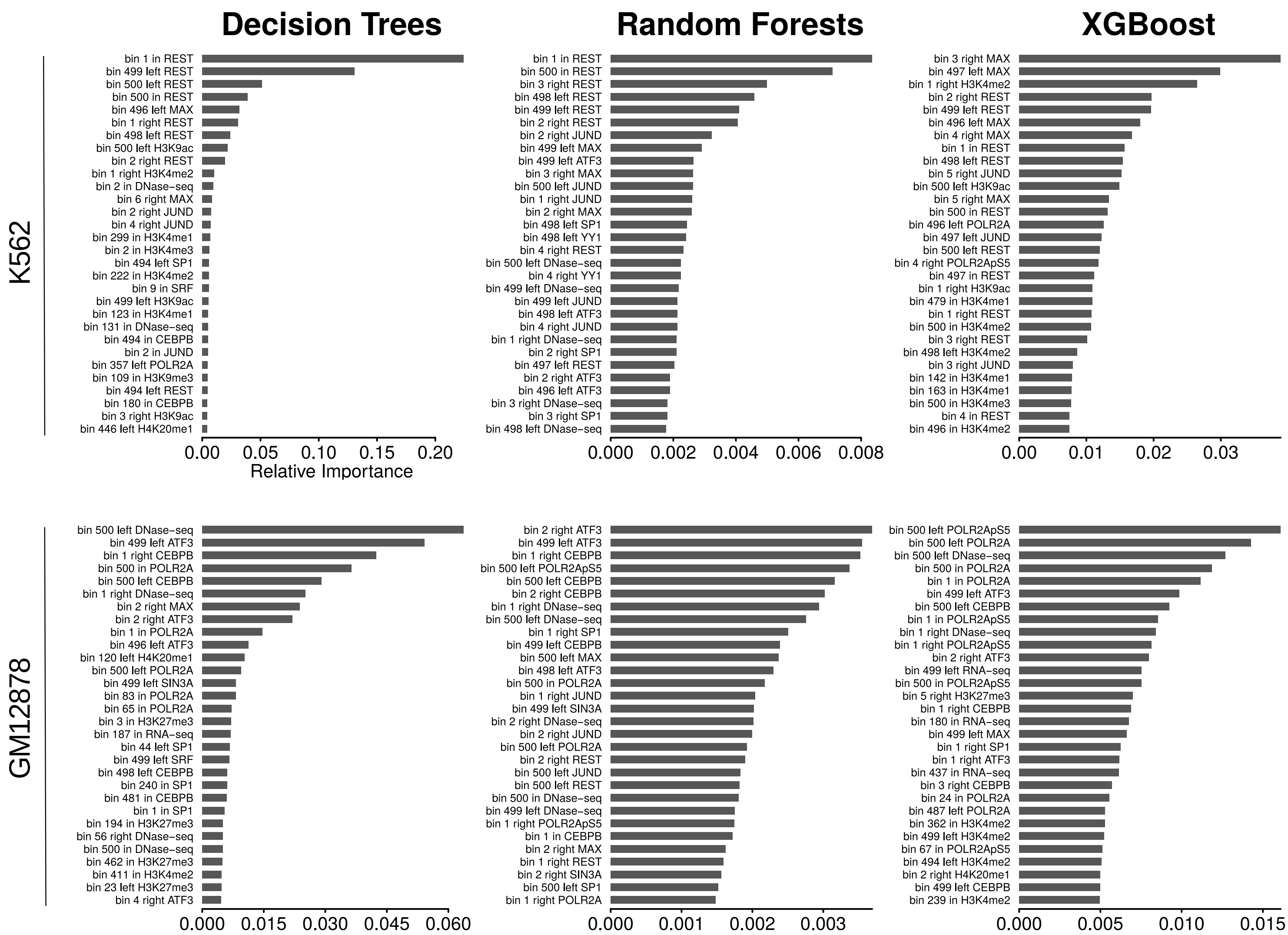

Decision Trees
Random Forests
XGBoost
K562
GM12878
Relative Importance
bin 1 in REST
bin 499 left REST
bin 500 left REST
bin 500 in REST
bin 496 left MAX
bin 1 right REST
bin 498 left REST
bin 500 left H3K9ac
bin 2 right REST
bin 1 right H3K4me2
bin 2 in DNase-seq
bin 6 right MAX
bin 2 right JUND
bin 4 right JUND
bin 299 in H3K4me1
bin 2 in H3K4me3
bin 494 left SP1
bin 222 in H3K4me2
bin 9 in SRF
bin 499 left H3K9ac
bin 123 in H3K4me1
bin 131 in DNase-seq
bin 494 in CEBPB
bin 2 in JUND
bin 357 left POLR2A
bin 109 in H3K9me3
bin 494 left REST
bin 180 in CEBPB
bin 3 right H3K9ac
bin 446 left H4K20me1
0.00 0.05 0.10 0.15 0.20
bin 1 in REST
bin 500 in REST
bin 3 right REST
bin 498 left REST
bin 499 left REST
bin 2 right REST
bin 2 right JUND
bin 499 left MAX
bin 499 left ATF3
bin 3 right MAX
bin 500 left JUND
bin 1 right JUND
bin 2 right MAX
bin 498 left SP1
bin 498 left YY1
bin 4 right REST
bin 500 left DNase-seq
bin 4 right YY1
bin 499 left DNase-seq
bin 499 left JUND
bin 498 left ATF3
bin 4 right JUND
bin 1 right DNase-seq
bin 2 right SP1
bin 497 left REST
bin 2 right ATF3
bin 496 left ATF3
bin 3 right DNase-seq
bin 3 right SP1
bin 498 left DNase-seq
0.000 0.002 0.004 0.006 0.008
bin 3 right MAX
bin 497 left MAX
bin 1 right H3K4me2
bin 2 right REST
bin 499 left REST
bin 496 left MAX
bin 4 right MAX
bin 1 in REST
bin 498 left REST
bin 5 right JUND
bin 500 left H3K9ac
bin 5 right MAX
bin 500 in REST
bin 496 left POLR2A
bin 497 left JUND
bin 500 left REST
bin 4 right POLR2ApS5
bin 497 in REST
bin 1 right H3K9ac
bin 479 in H3K4me1
bin 1 right REST
bin 500 in H3K4me2
bin 3 right REST
bin 498 left H3K4me2
bin 3 right JUND
bin 142 in H3K4me1
bin 163 in H3K4me1
bin 500 in H3K4me3
bin 4 in REST
bin 496 in H3K4me2
0.00 0.01 0.02 0.03
bin 500 left DNase-seq
bin 499 left ATF3
bin 1 right CEBPB
bin 500 in POLR2A
bin 500 left CEBPB
bin 1 right DNase-seq
bin 2 right MAX
bin 2 right ATF3
bin 1 in POLR2A
bin 496 left ATF3
bin 120 left H4K20me1
bin 500 left POLR2A
bin 499 left SIN3A
bin 83 in POLR2A
bin 65 in POLR2A
bin 3 in H3K27me3
bin 187 in RNA-seq
bin 44 left SP1
bin 499 left SRF
bin 498 left CEBPB
bin 240 in SP1
bin 481 in CEBPB
bin 1 in SP1
bin 194 in H3K27me3
bin 56 right DNase-seq
bin 500 in DNase-seq
bin 462 in H3K27me3
bin 411 in H3K4me2
bin 23 left H3K27me3
bin 4 right ATF3
0.000 0.015 0.030 0.045 0.060
bin 2 right ATF3
bin 499 left ATF3
bin 1 right CEBPB
bin 500 left POLR2ApS5
bin 500 left CEBPB
bin 2 right CEBPB
bin 1 right DNase-seq
bin 500 left DNase-seq
bin 1 right SP1
bin 499 left CEBPB
bin 500 left MAX
bin 498 left ATF3
bin 500 in POLR2A
bin 1 right JUND
bin 499 left SIN3A
bin 2 right DNase-seq
bin 2 right JUND
bin 500 left POLR2A
bin 2 right REST
bin 500 left JUND
bin 500 left REST
bin 500 in DNase-seq
bin 499 left DNase-seq
bin 1 right POLR2ApS5
bin 1 in CEBPB
bin 2 right MAX
bin 1 right REST
bin 2 right SIN3A
bin 500 left SP1
bin 1 right POLR2A
0.000 0.001 0.002 0.003
bin 500 left POLR2ApS5
bin 500 left POLR2A
bin 500 left DNase-seq
bin 500 in POLR2A
bin 1 in POLR2A
bin 499 left ATF3
bin 500 left CEBPB
bin 1 in POLR2ApS5
bin 1 right DNase-seq
bin 1 right POLR2ApS5
bin 2 right ATF3
bin 499 left RNA-seq
bin 500 in POLR2ApS5
bin 5 right H3K27me3
bin 1 right CEBPB
bin 180 in RNA-seq
bin 499 left MAX
bin 1 right SP1
bin 1 right ATF3
bin 437 in RNA-seq
bin 3 right CEBPB
bin 24 in POLR2A
bin 487 left POLR2A
bin 362 in H3K4me2
bin 499 left H3K4me2
bin 67 in POLR2ApS5
bin 494 left H3K4me2
bin 2 right H4K20me1
bin 499 left CEBPB
bin 239 in H3K4me2
0.000 0.005 0.010 0.015

### Figure S5

#### **A)** Decision Trees (K562)

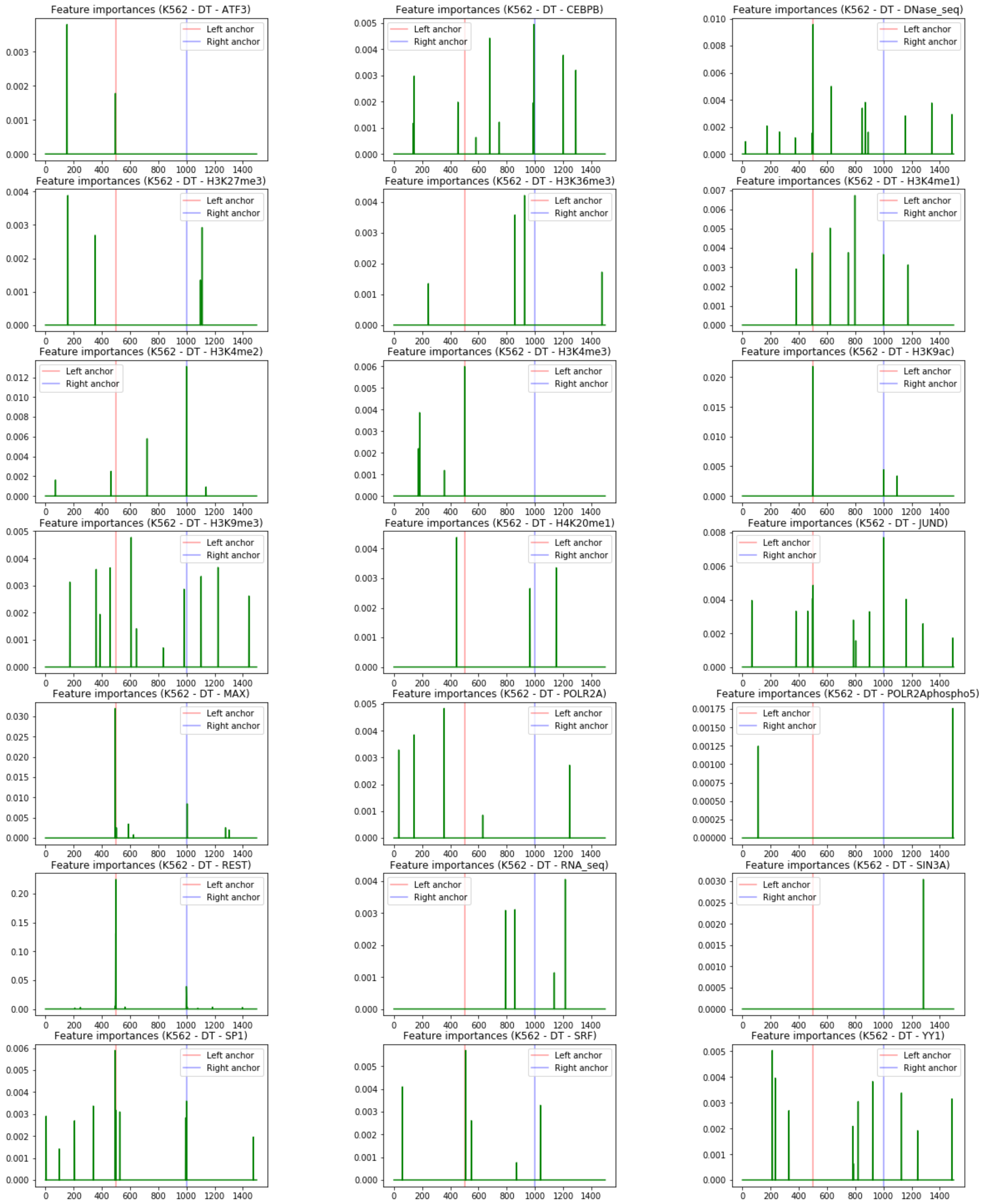

**B)** Random Forests (K562)

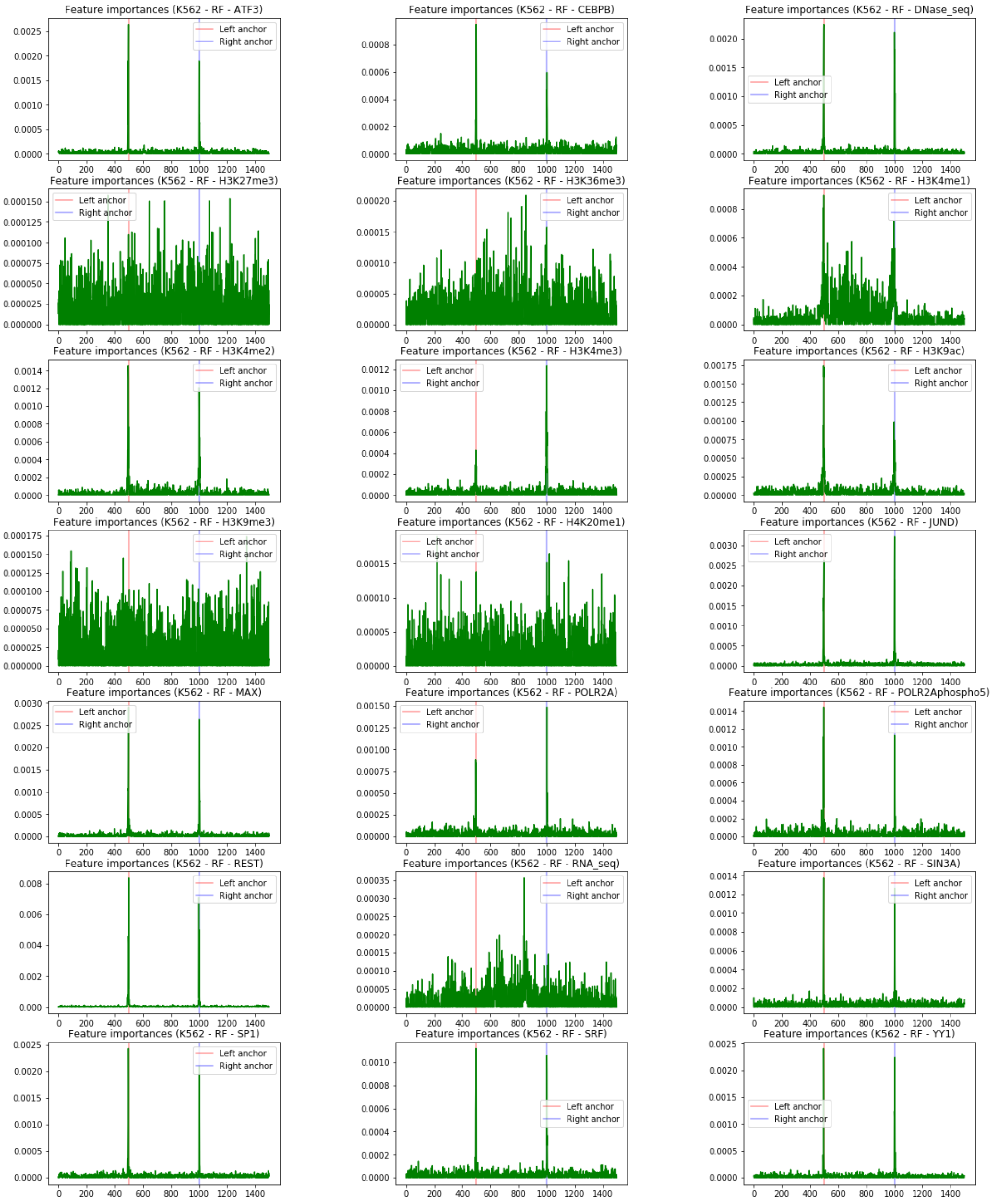

#### **C)** XGBoost (K562)

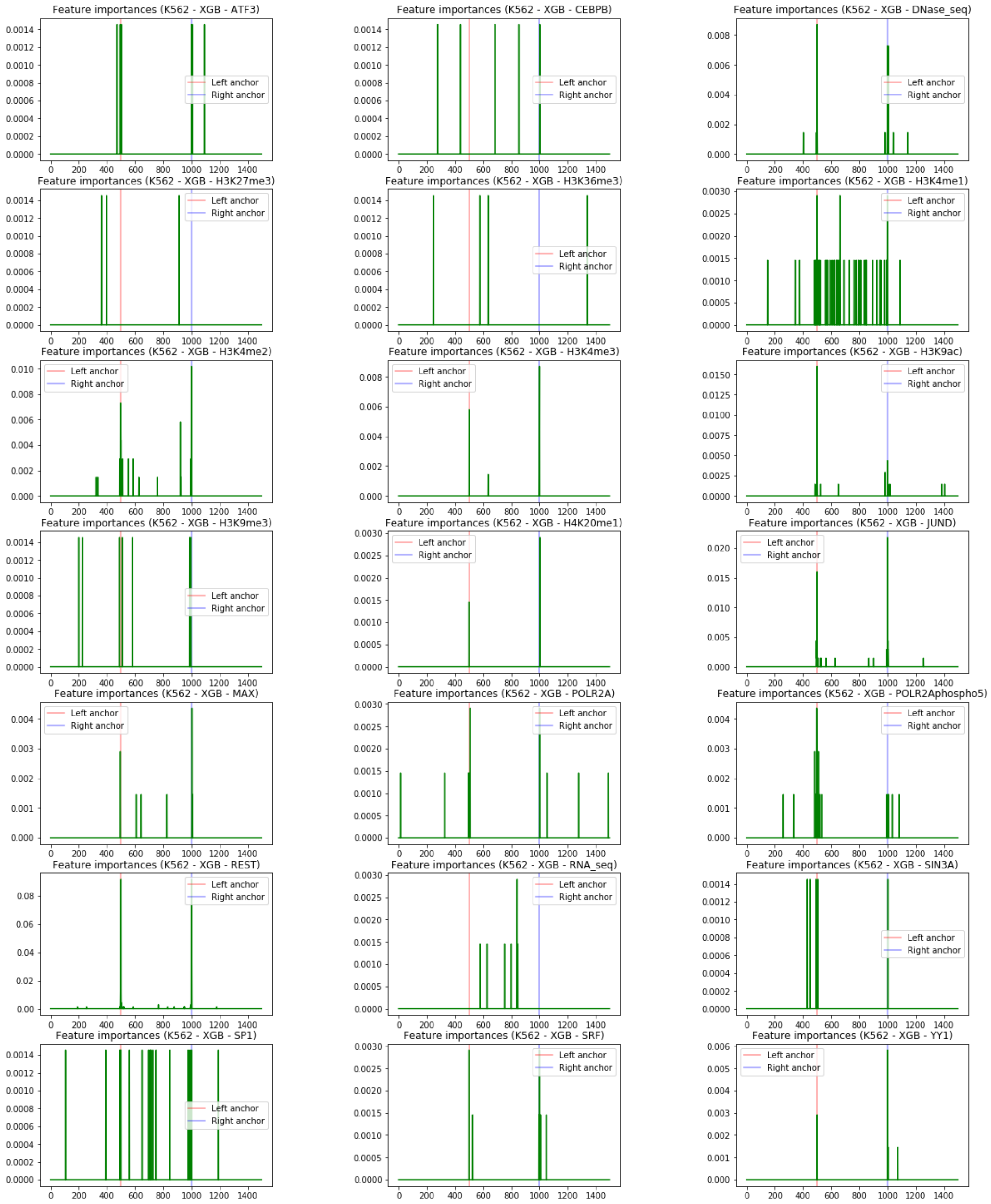

#### **D)** Decision Trees (GM12878)

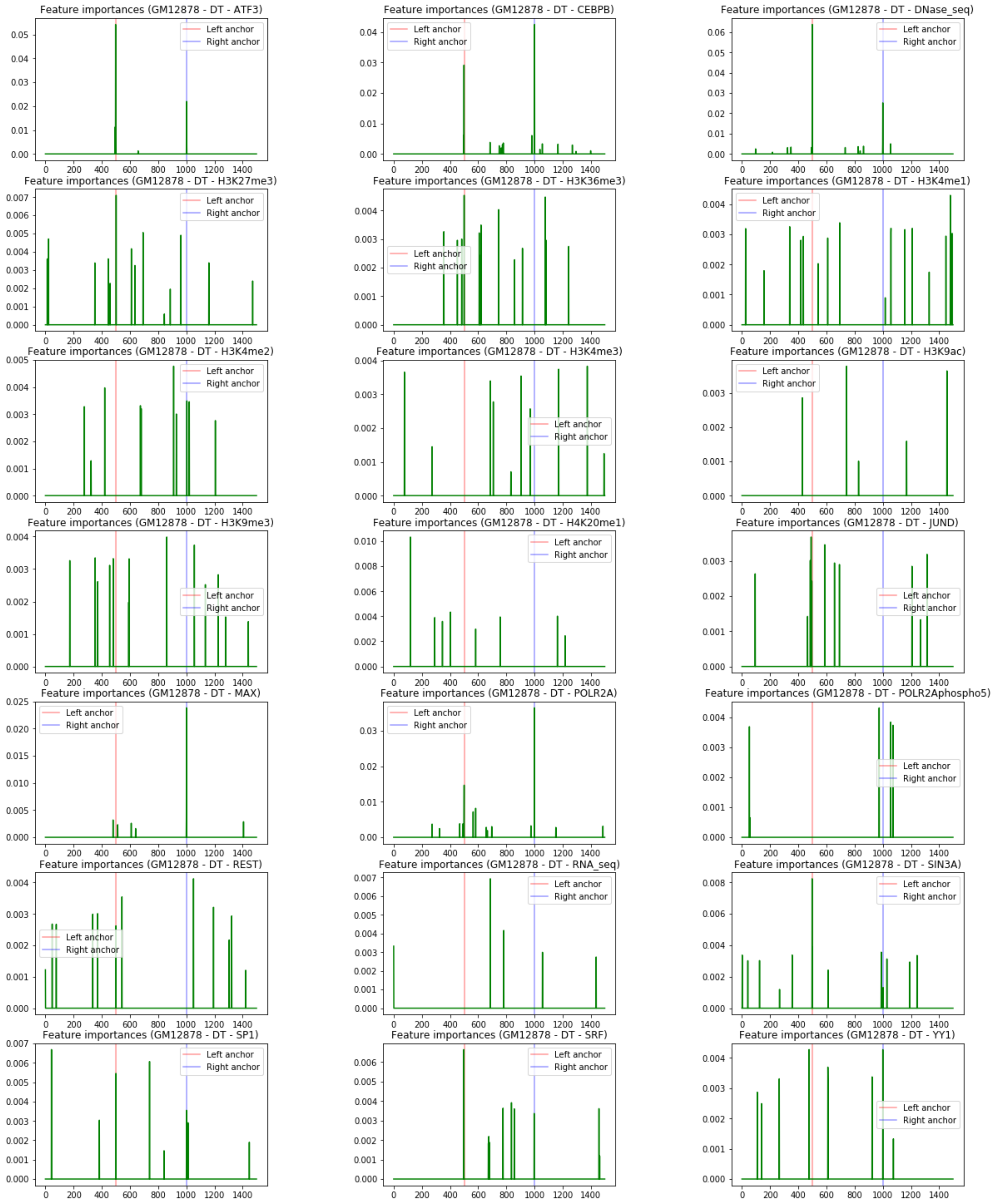

**E)** Random Forests (GM12878)

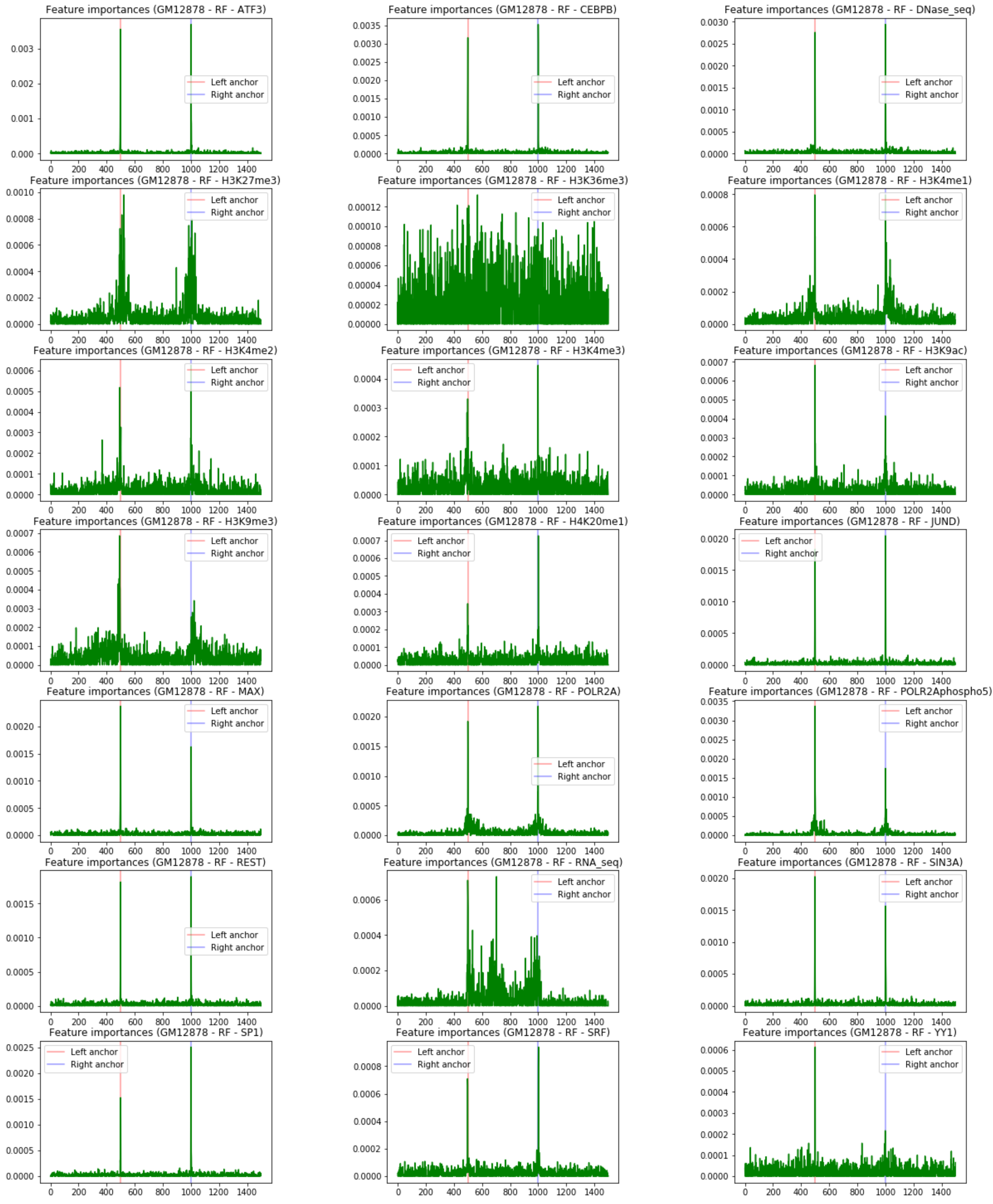

**F)** XGBoost (GM12878)

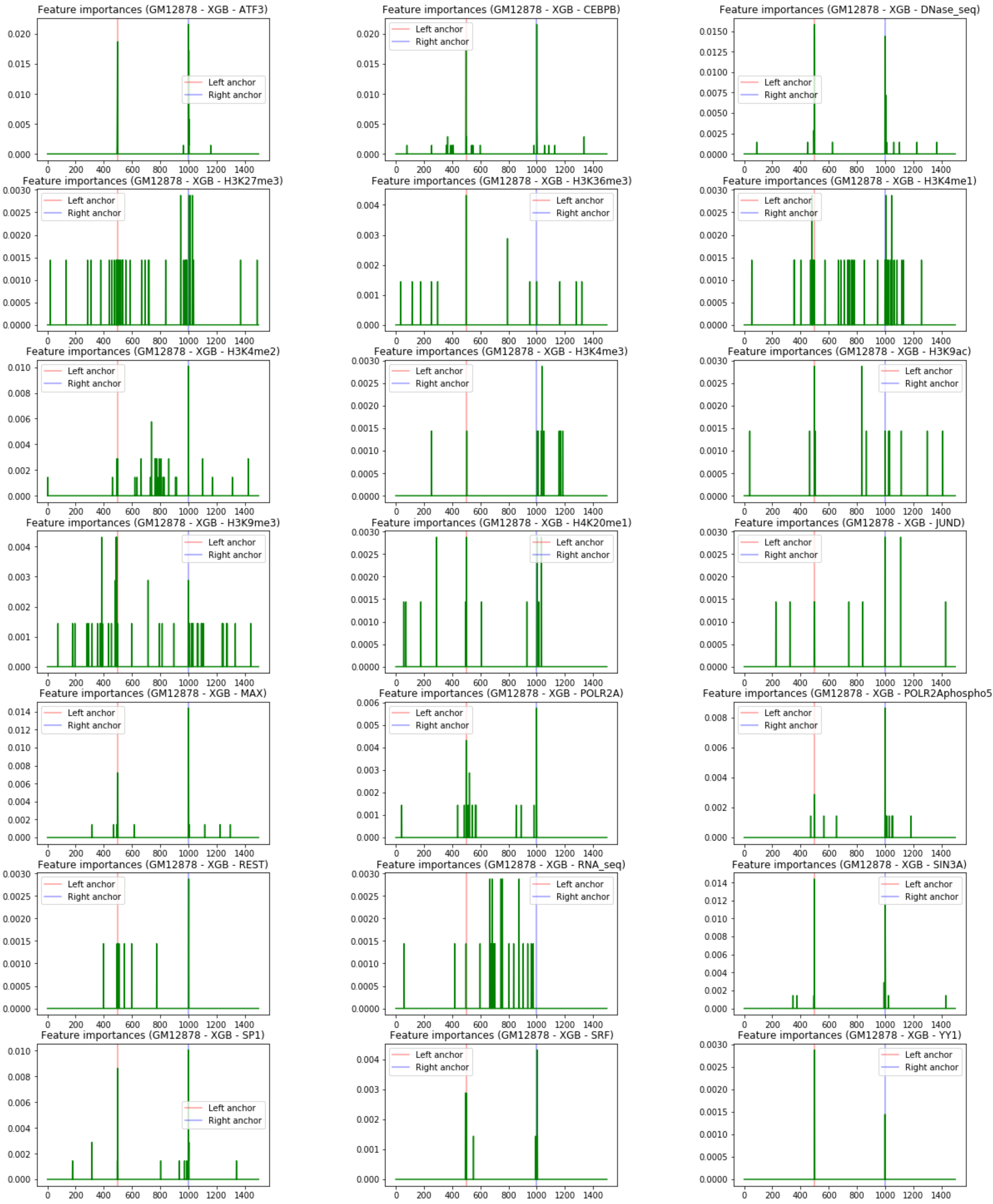

### Figure S6

#### A) Architectural

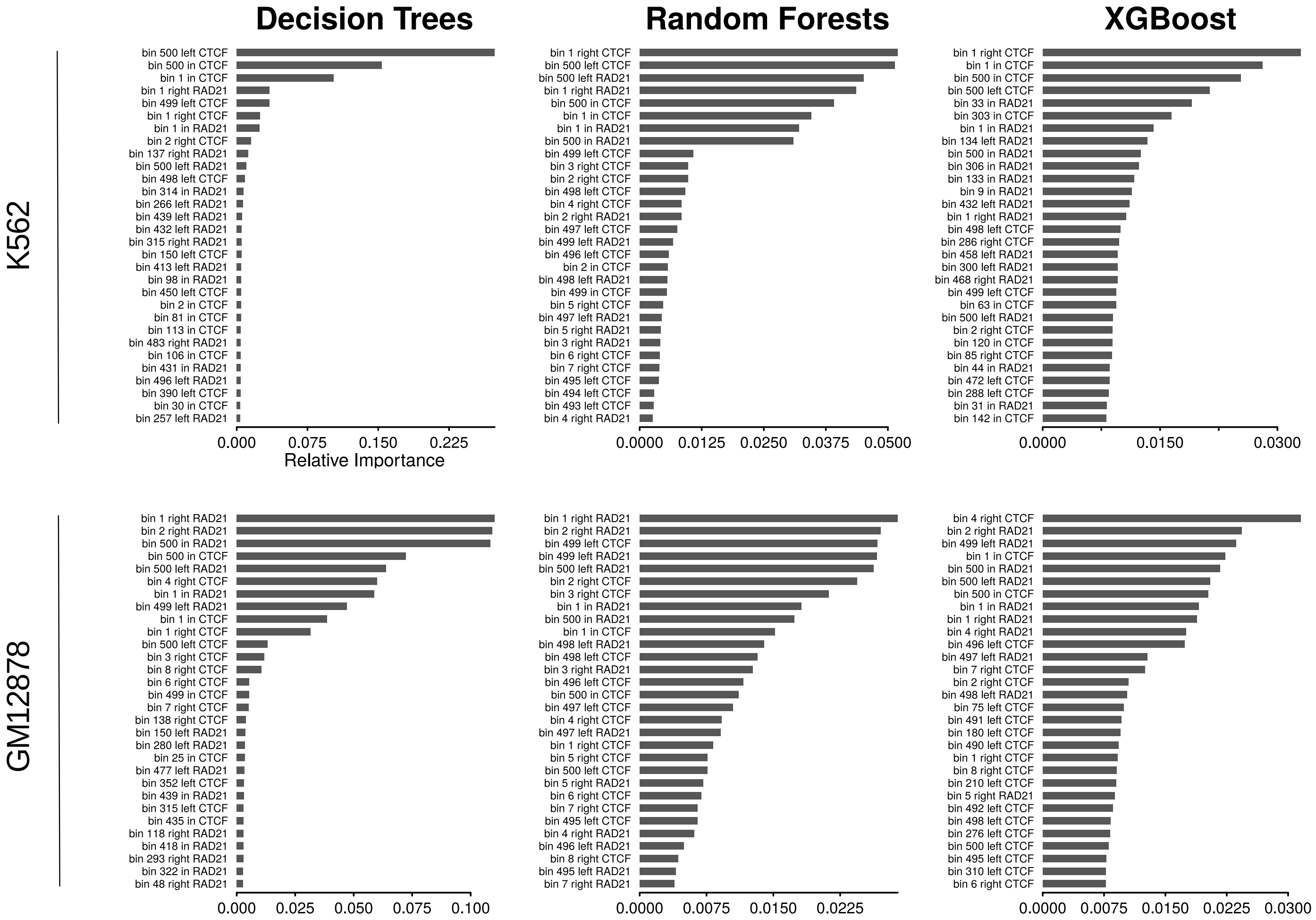


#### B) Transcription factors

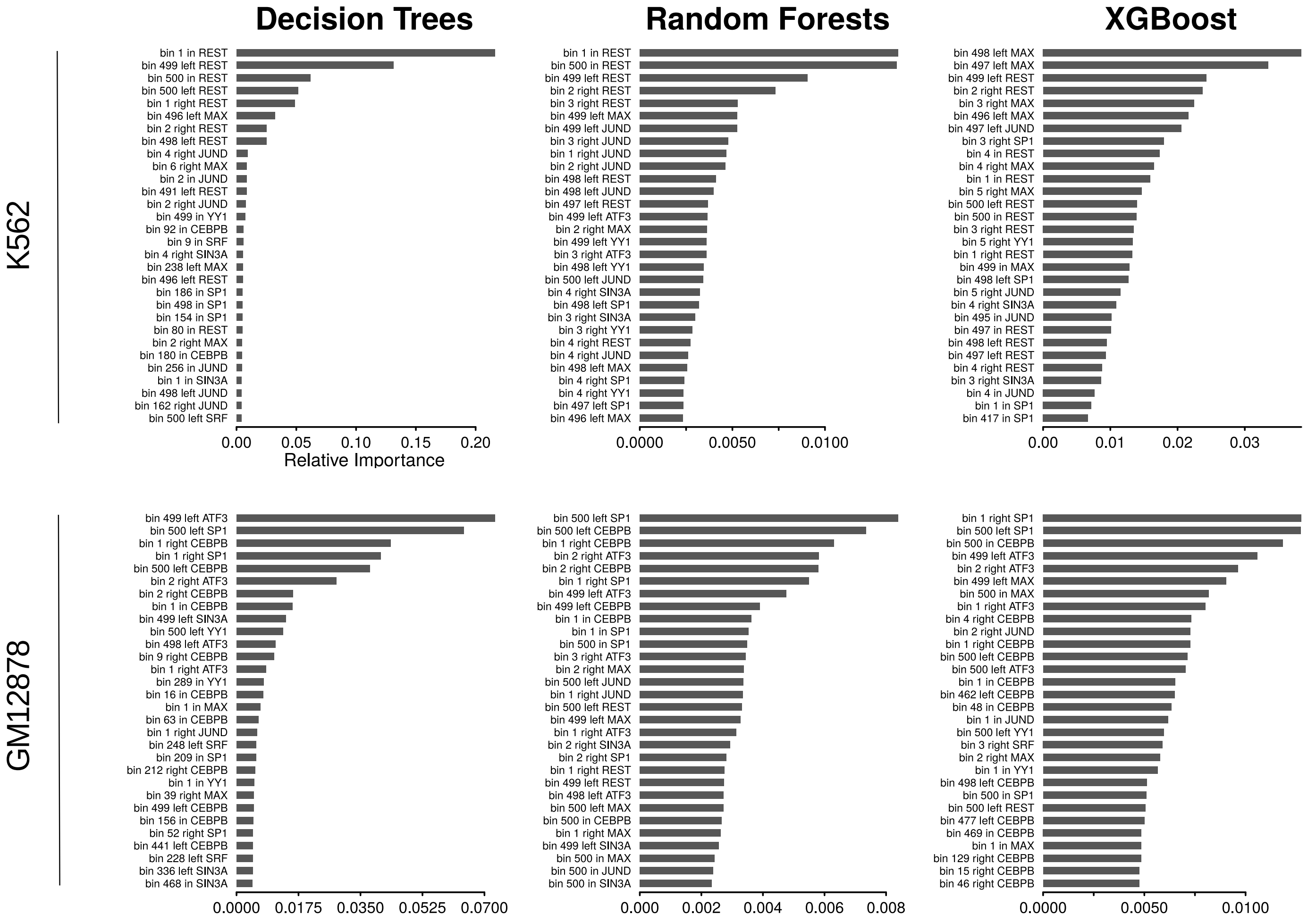

#### **C)** Architectural-anchors

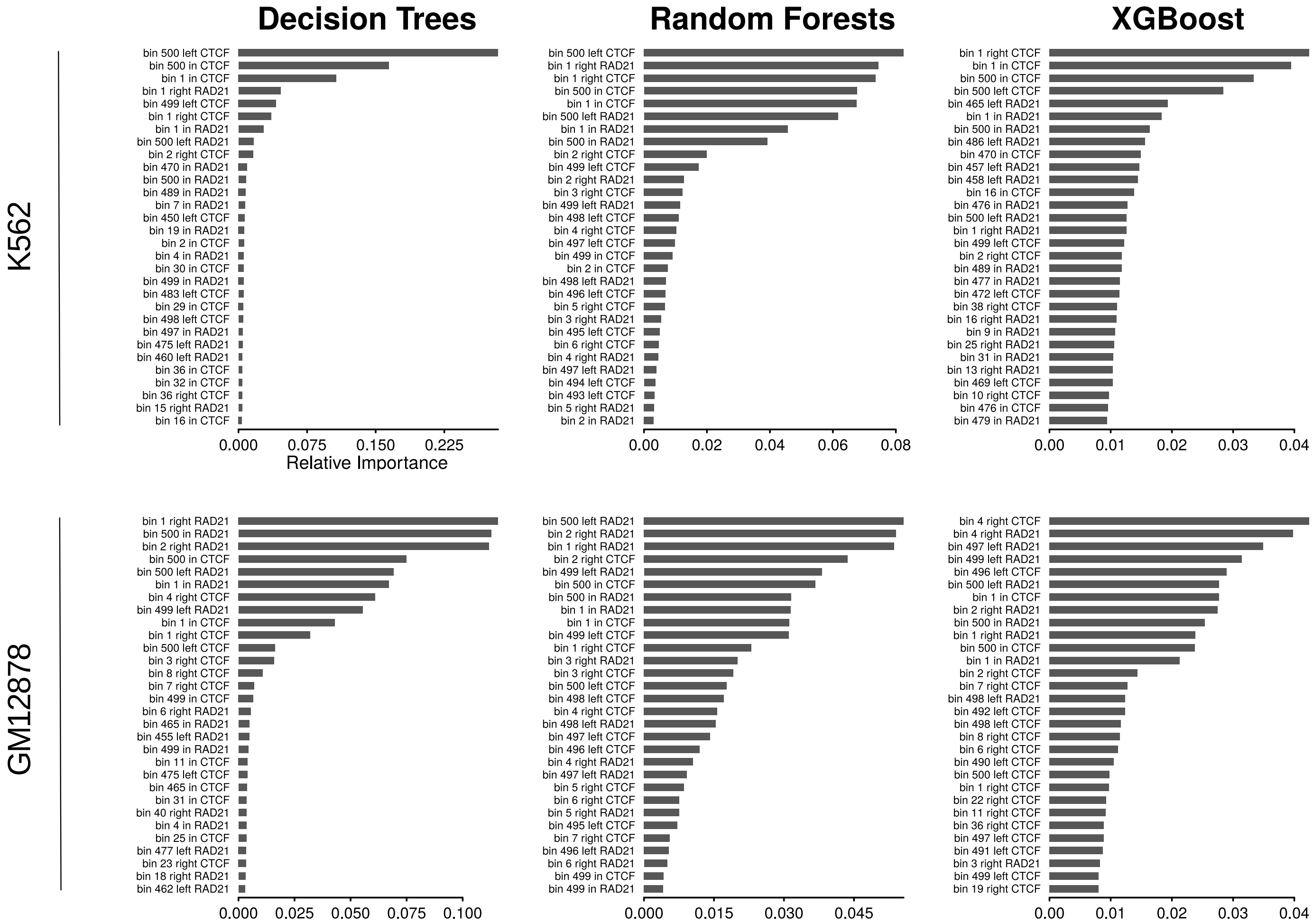


#### **D)** Transcription factors-anchors

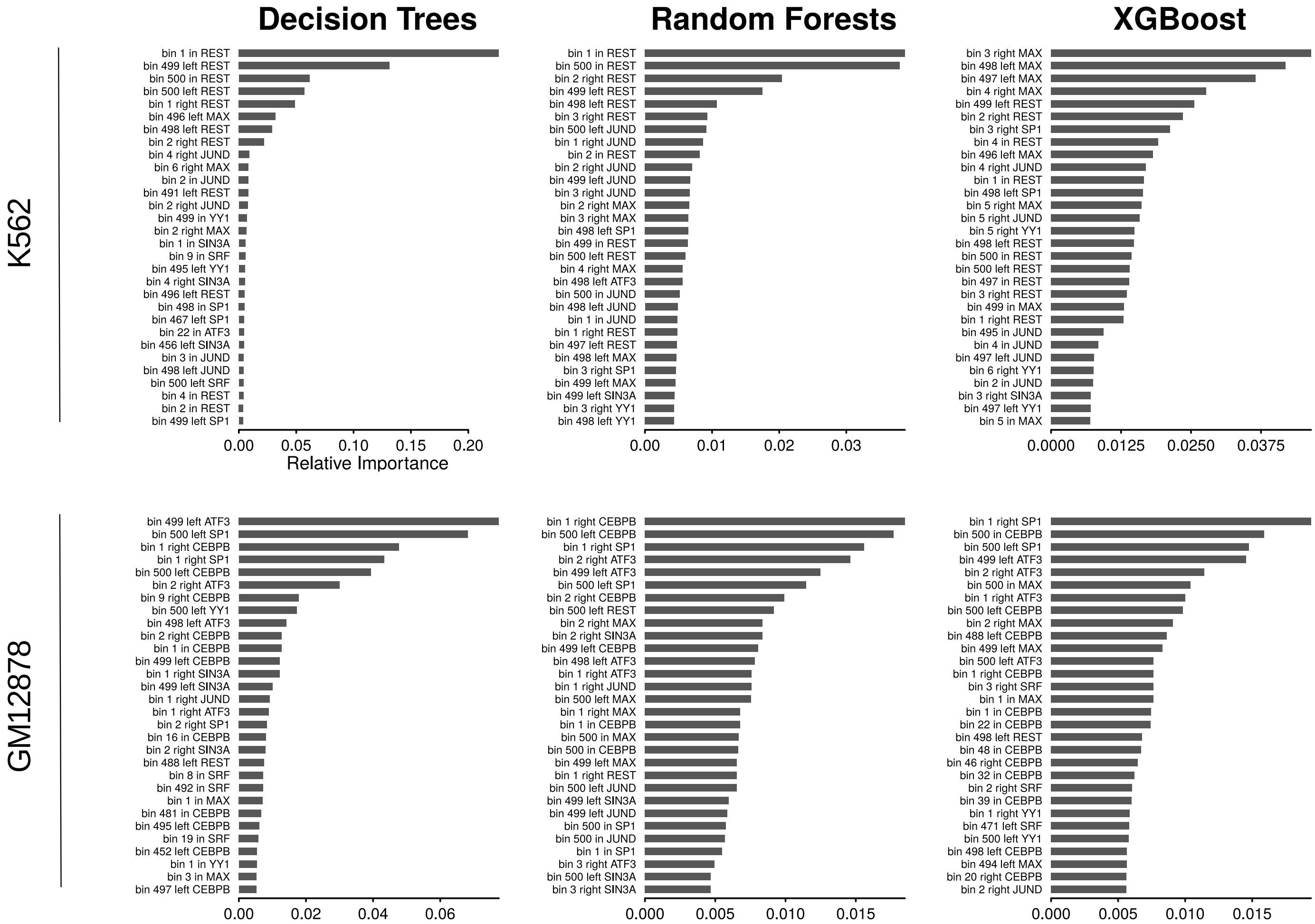

**Table S1.** Public data used in this study.

| Accession | Assay | Cell line | Target | Source |
|---|---|---|---|---|
| ENCSR752QCX | ChIA-PET | GM12878 | RAD21 | ENCODE |
| GSE18199 | Hi-C | GM12878 | - | GEO |
| ENCSR000COR | RNA-seq | GM12878 | - | ENCODE |
| ENCSR000EMT | DNase-seq | GM12878 | - | ENCODE |
| ENCSR000BJY | ChIP-seq | GM12878 | ATF3 | ENCODE |
| ENCSR000BRX | ChIP-seq | GM12878 | CEBPB | ENCODE |
| ENCSR000DKV | ChIP-seq | GM12878 | CTCF | ENCODE |
| ENCSR000DRX | ChIP-seq | GM12878 | H3K27me3 | ENCODE |
| ENCSR000DRW | ChIP-seq | GM12878 | H3K36me3 | ENCODE |
| ENCSR000AKF | ChIP-seq | GM12878 | H3K4me1 | ENCODE |
| ENCSR000AKG | ChIP-seq | GM12878 | H3K4me2 | ENCODE |
| ENCSR057BWO | ChIP-seq | GM12878 | H3K4me3 | ENCODE |
| ENCSR000AKH | ChIP-seq | GM12878 | H3K9ac | ENCODE |
| ENCSR000AOX | ChIP-seq | GM12878 | H3K9me3 | ENCODE |
| ENCSR000AKI | ChIP-seq | GM12878 | H4K20me1 | ENCODE |
| ENCSR000DYS | ChIP-seq | GM12878 | JUND | ENCODE |
| ENCSR000DZF | ChIP-seq | GM12878 | MAX | ENCODE |
| ENCSR000EAD | ChIP-seq | GM12878 | POLR2A | ENCODE |
| ENCSR000BIF | ChIP-seq | GM12878 | POLR2AphosphoS5 | ENCODE |
| ENCSR000EAC | ChIP-seq | GM12878 | RAD21 | ENCODE |
| ENCSR000BQS | ChIP-seq | GM12878 | REST | ENCODE |
| ENCSR000DYX | ChIP-seq | GM12878 | SIN3A | ENCODE |
| ENCSR000BHK | ChIP-seq | GM12878 | SP1 | ENCODE |
| ENCSR000BGE | ChIP-seq | GM12878 | SRF | ENCODE |
| ENCSR000EUM | ChIP-seq | GM12878 | YY1 | ENCODE |
| ENCSR000FDB | ChIA-PET | K562 | RAD21 | ENCODE |
| GSE18199 | Hi-C | K562 | - | GEO |
| ENCSR000COK | RNA-seq | K562 | - | ENCODE |
| ENCSR000EPC | DNase-seq | K562 | - | ENCODE |
| ENCSR000BNU | ChIP-seq | K562 | ATF3 | ENCODE |
| ENCSR000EHE | ChIP-seq | K562 | CEBPB | ENCODE |
| ENCSR000DMA | ChIP-seq | K562 | CTCF | ENCODE |

| | | | | |
|---|---|---|---|---|
| ENCSR000EWB | ChIP-seq | K562 | H3K27me3 | ENCODE |
| ENCSR000DWB | ChIP-seq | K562 | H3K36me3 | ENCODE |
| ENCSR000EWC | ChIP-seq | K562 | H3K4me1 | ENCODE |
| ENCSR000AKT | ChIP-seq | K562 | H3K4me2 | ENCODE |
| ENCSR668LDD | ChIP-seq | K562 | H3K4me3 | ENCODE |
| ENCSR000EVZ | ChIP-seq | K562 | H3K9ac | ENCODE |
| ENCSR000APE | ChIP-seq | K562 | H3K9me3 | ENCODE |
| ENCSR000AKX | ChIP-seq | K562 | H4K20me1 | ENCODE |
| ENCSR000EGN | ChIP-seq | K562 | JUND | ENCODE |
| ENCSR000BLP | ChIP-seq | K562 | MAX | ENCODE |
| ENCSR000BMR | ChIP-seq | K562 | POLR2A | ENCODE |
| ENCSR000BKR | ChIP-seq | K562 | POLR2AphosphoS5 | ENCODE |
| ENCSR000FAD | ChIP-seq | K562 | RAD21 | ENCODE |
| ENCSR137ZMQ | ChIP-seq | K562 | REST | ENCODE |
| ENCSR000BLR | ChIP-seq | K562 | SIN3A | ENCODE |
| ENCSR991ELG | ChIP-seq | K562 | SP1 | ENCODE |
| ENCSR000BLK | ChIP-seq | K562 | SRF | ENCODE |
| ENCSR000BMH | ChIP-seq | K562 | YY1 | ENCODE |

**Table S2.** Machine learning performance of models trained with architectural factors binding information. K562 cell line.

| Algorithm | Accuracy | Precision | Recall | F1-Score |
|---|---|---|---|---|
| **Decision Trees** | 0.8546 | 0.8547 | 0.8546 | 0.8546 |
| **Random Forests** | 0.9124 | 0.9201 | 0.9125 | 0.9121 |
| **XGBoost** | 0.9459 | 0.9470 | 0.9460 | 0.9459 |

**Table S3.** Machine learning performance of models trained with architectural factors binding information. GM12878 cell line.

| Algorithm | Accuracy | Precision | Recall | F1-Score |
|---|---|---|---|---|
| **Decision Trees** | 0.8437 | 0.8442 | 0.8438 | 0.8437 |
| **Random Forests** | 0.8866 | 0.8882 | 0.8866 | 0.8865 |
| **XGBoost** | 0.9299 | 0.9312 | 0.9300 | 0.9299 |

**Table S4.** Machine learning performance of models trained with transcription factors binding information. K562 cell line.

| Algorithm | Accuracy | Precision | Recall | F1-Score |
|---|---|---|---|---|
| **Decision Trees** | 0.8386 | 0.8387 | 0.8387 | 0.8387 |
| **Random Forests** | 0.7709 | 0.7756 | 0.7709 | 0.7698 |
| **XGBoost** | 0.9360 | 0.9367 | 0.9361 | 0.9360 |

**Table S5.** Machine learning performance of models trained with transcription factors binding information. GM12878 cell line.

| Algorithm | Accuracy | Precision | Recall | F1-Score |
|---|---|---|---|---|
| **Decision Trees** | 0.6612 | 0.6644 | 0.6613 | 0.6603 |
| **Random Forests** | 0.7769 | 0.7877 | 0.7770 | 0.7753 |
| **XGBoost** | 0.7917 | 0.7937 | 0.7917 | 0.7915 |

**Table S6.** Machine learning performance of models trained with architectural factors binding information at loop anchors. K562 cell line.

| Algorithm | Accuracy | Precision | Recall | F1-Score |
|---|---|---|---|---|
| **Decision Trees** | 0.8736 | 0.8760 | 0.8737 | 0.8734 |
| **Random Forests** | 0.9254 | 0.9290 | 0.9254 | 0.9252 |
| **XGBoost** | 0.9421 | 0.9430 | 0.9422 | 0.9421 |

**Table S7.** Machine learning performance of models trained with architectural factors binding information at loop anchors. GM12878 cell line.

| Algorithm | Accuracy | Precision | Recall | F1-Score |
|---|---|---|---|---|
| **Decision Trees** | 0.8520 | 0.8528 | 0.8521 | 0.8519 |
| **Random Forests** | 0.9050 | 0.9086 | 0.9051 | 0.9048 |
| **XGBoost** | 0.9290 | 0.9300 | 0.9290 | 0.9290 |

**Table S8.** Machine learning performance of models trained with transcription factors binding information at loop anchors. K562 cell line.

| Algorithm | Accuracy | Precision | Recall | F1-Score |
|---|---|---|---|---|
| **Decision Trees** | 0.8508 | 0.8509 | 0.8508 | 0.8508 |
| **Random Forests** | 0.8417 | 0.8463 | 0.8417 | 0.8411 |
| **XGBoost** | 0.9345 | 0.9350 | 0.9346 | 0.9345 |

**Table S9.** Machine learning performance of models trained with transcription factors binding information at loop anchors. GM12878 cell line.

| Algorithm | Accuracy | Precision | Recall | F1-Score |
|---|---|---|---|---|
| **Decision Trees** | 0.6801 | 0.6810 | 0.6802 | 0.6801 |
| **Random Forests** | 0.7852 | 0.7917 | 0.7853 | 0.7844 |
| **XGBoost** | 0.7903 | 0.7930 | 0.7903 | 0.7900 |
